# Unveiling the genetic basis of the low pH response in the acidophilic yeast *Maudiozyma bulderi* as a potential host for biorefinery

**DOI:** 10.1101/2025.06.27.661920

**Authors:** Laura Natalia Balarezo-Cisneros, Alistair Hanak, Leo Zeef, Aleksandr Mironov, Fernando Valle, Daniela Delneri

## Abstract

Non-conventional yeasts represent a great genetic and phenotypic diversity with potential for industrial strain development in the bio-production of green chemicals. In recent years, mass genome sequencing of non-conventional yeasts has opened avenues to improved understanding of transcriptional networks and phenotypic plasticity and gene function, including the discovery of novel genes. Here, we investigated the expressional and morphological changes at low-pH in three strains of the acidophilic yeast *Maudiozyma bulderi* (previously *Kazachstania bulderi* and *Saccharomyces bulderi*): CBS 8638, CBS 8639 and NRRL Y-27205. The comparison of the transcriptome of cells growing in a bioreactor at pH=5.5 vs pH= 2.5, primarily showed dysregulation of genes involved in cell wall integrity, with NRRL Y-27205 the least acidophilic strain, showing the largest transcriptional response when compared to the other two strains. We identified four uncharacterised genes, unique to *M. bulderi,* and predicted function as transporters, upregulated at low pH. Microscopy studies showed that *M. bulderi* cell wall is not damaged in acidic environment, and the membrane lipid composition remains stable at low pH, unlike *S. cerevisiae*. Overall, our data on transcriptional variability in *M. bulderi* highlights genes and cellular pathways involved in the acidophilic adaptation of this species and can aid further strain development.

## 1 Introduction

Microorganisms are widely used as cell factories in biorefineries, where maintaining optimal conditions for production host species accounts for significant operational costs (Kamm & Kamm, 2004; Abdel-Rahman *et al*., 2011;Sarkar *et al*., 2012; Prado-Rubio *et al*., 2020). Organic acids, such as lactate, pyruvate, citrate, gluconate, and itaconate, are important platform chemicals used to produce a variety of commercial products, including biodegradable polymers and numerous sustainable chemical compounds (Sauer *et al.,* 2008; de Oliveira *et al.,* 2018). Because of this, several technologies have been developed to produce organic acids by fermentation of microorganisms. However, most production microorganisms in the industry, require a neutral pH for optimal performance during fermentation. As the organic acids are synthesized and accumulate in the culture media, the pH decreases, necessitating the addition of a base to neutralize the organic acid, resulting in the formation of the organic acid salt. For subsequent applications, the acid form of the organic acids is typically required, meaning that the neutralizing cation must be removed, disposed of, or recycled. This significantly increases production costs (Cao *et al.,* 2002; Chen & Nielsen, 2016; Ahmad *et al*., 2020). Because of this, there is a need to develop new production microorganism capable of producing efficiently organic acids at low pH. Furthermore, it is also desirable to understand the mechanisms that acidophilic microorganisms use to thrive at low pH, to try to transfer this capacity to more traditional production hosts.

Non-conventional yeasts, including extremophilic species with natural adaptations to hostile environments, possess great genetic and phenotypic diversity that is yet untapped.These yeasts are increasingly viewed as an important resource in mitigating the need for expensive remediation measures during fermentations (Mukherjee *et al*., 2017; Rebello *et al*., 2018; Yamakawa *et al*., 2020; Geijer *et al*., 2022). In recent years the short-read genome sequencing of many non-conventional yeast species has accelerated (Hittinger *et al.,* 2015; Shen *et al*., 2018; Opulente *et al.,* 2024). Despite these advances, many species still lack high quality genome assemblies and annotation which are required to pinpoint specific mechanisms of stress tolerance.

*Maudiozyma bulderi* (previously known as *Saccharomyces bulderi* and *Kazachstania bulderi)* is a remarkable non-conventional yeast, first isolated from maize silage (Middelhoven *et al*., 2000), known to thrive in low pH conditions. It is capable of maintaining a high growth rate at pH between 2.5 and 5 (van Dijken *et al*., 2002), and high concentrations of several organic acids (Balarezo-Cisneros *et al*., 2023). A high-quality genome of *M. bulderi*, including structural and functional annotation, has recently been published (Balarezo-Cisneros *et al*., 2023) allowing further investigation of the genetic underpinning of acidophilic traits.

The response to low pH in *S. cerevisiae* is well studied, as are the mechanisms employed to maintain a stable cytosolic pH. To work efficiently in an acidic environment, cells must maintain a proton gradient over the plasma membrane for ATP synthesis and nutrient uptake (Nishimura, *et al.,* 1998; Mongan and Case, 2005; Orij *et al*., 2011) The yeast plasma membrane is impermeable to H+ ions and contains membrane bound H+ ATPases, primarily Pma1p, which are responsible for proton efflux, pumping H+ ions out of the cell at the cost of ATP (Serrano *et al.,* 1986; Goossens *et al.,* 2000). Low pH can also cause damage to cell wall structures and disrupt cell integrity (Simpson and Hammond, 1989; Graff *et al.,* 2008; de Lucena *et al*., 2015). Maintenance of the cell wall is regulated primarily via the cell wall integrity (CWI) pathway (Levin, 2011), which is crucial for regulation of the yeast response to low pH (de Lucena *et al.,* 2012; de Lucena *et al.,* 2015). The CWI regulates the repair of the cell wall caused by environmental stresses, via several membrane bound sensor proteins that detect stressors and trigger phosphorylation cascades. The main sensor responsible for low pH detection is Mid2p which recruits the guanine nucleotide exchange proteins Rom1p and Rom2p to the cell membrane where they activate the GTPase Rho1p.

Rho1p then triggers a cascade resulting in the activation of cell wall biogenesis genes including *FKS1* and *FKS2* (Claret *et al.,* 2005). Low pH can also induce other stress related pathways such as the high osmolarity glycerol (HOG) pathway and general stress response (GSR), which include several overlapping genes (Kapteyn *et al*., 2001; Kawahata *et al*., 2006; Zakrzewska *et al.,* 2007; de Melo *et al.,* 2010). Cell pH homeostasis is supported by auxiliary systems including the vacuole amongst other organelles. The vacuole is kept at a more acidic pH relative to the cytosol via the action of H+ ATPases including Vma1p, which transports H+ inside the vacuole (Martínez-Muñoz, & Kane, 2008). Additionally, cation antiporter proteins including Nha1p, Vcx1p and Vnx1p are involved in the movement of H+ ions across the plasma membrane and organellar membranes contributing to removal of excess H+ ions in the cytosol (fPrior *et al.,* 1996; Sychrová *etal.,* 1999). The structure and composition of the cell membrane can influence pH tolerance, with changes impacting membrane fluidity (Klose *et al.,* 2012; Santos & Preta, 2018; Ferraz *et al.,* 2021). The yeast cell membrane consists primarily of phospholipids, sphingolipids, sterol lipids. Sterol lipids such as ergosterol are the main candidates for response to low pH stress in *S. cerevisiae* (Vanegas *et al.,* 2012). Ergosterol impacts the permeability of the cell membrane, this is particularly important in the response to organic acids as undissociated forms are soluble in the lipid bilayer (Ferraz *et al.,* 2023). Undissociated organic acids can pass into the cell where they dissociate in the higher pH of the cytosol acidifying it. Ergosterol is also necessary for the formation of lipid rafts and the localisation of membrane-bound transporters and glycosylphosphatidylinositol anchored proteins involved in stress detection (Eisenkolb *et al.,* 2002). It has also been shown to be necessary to localise H+ ATPases including Pma1p to the cell membrane in *Candida albicans* (Urbanek *et al.,* 2022). Increasing ergosterol content of cell membranes in *S. cerevisiae* has been shown to improve tolerance to specific inorganic acids (Guo *et al.,* 2018; Ferraz *et al.,* 2023).

In some non-conventional yeasts, the response to low pH has been studied. Species such as *Debaryomyces hansenii* and *Candida glabrata* show a similar response to *S. cerevisiae* in terms of lipid composition, where membrane fluidity was reduced by increased sterol lipids in response to strong inorganic acid and lactic acid (Turk *et al*., 2007 Lin *et al*., 2017; Ferraz *et al.,* 2023). Likewise, compared to *S. cerevisiae*)the low pH tolerant species *Zygosacchormyces parabilii* has been shown to express a similar but more extreme reduction in glucans and mannans in the cell wall (Berterame *et al*., 2016; Kuanyshev *et al*., 2016;, while *Pichia anomala* showed increased expression of membrane H+ ATPases and mitochondrial ATP producing genes in response to acid pH (Fletcher *et al*., 2015); and *Pichia kudriavzevii* upregulates genes associated with membrane transport, amino acid biosynthesis and metabolism of fatty acids and glycophospholipids (Ji *et al*., 2020). Understanding of the diverse responses to low pH adopted by different yeast species, could help targeted approaches to strain development, paving the way for the use of new species as industrial production hosts.

Here, we studied the cellular response to low pH by investigating the transcriptional landscape in *M. bulderi,* alongside cell morphology and lipid composition (Fig.1). We carried out fermentations with *M. bulderi* at pH maintained at either 5.5 or 2.5. RNA-seq analysis was carried out to identify expressional changes that are related to the high growth of *M. bulderi* at low pH. We found that a multifaceted response of *M. bulderi* to acidic environments with genes implicated in the GSR and CWI pathway, displaying increased expression at low pH. Comparative analysis between strains revealed distinct transcriptional responses, recapitulating phenotypic diversity among *M. bulderi* populations, and we identified DE species-specific genes. We showed that, unlike *S. cerevisiae*, *M. bulderi* cell wall is not damaged by low pH and the lipid composition also remains unchanged. Overall, our study sheds light on the pathways underpinning the remarkable tolerance of *M. bulderi* to acidic environments and will aid further engineering strategies to develop this species as production host.

**Figure 1.**
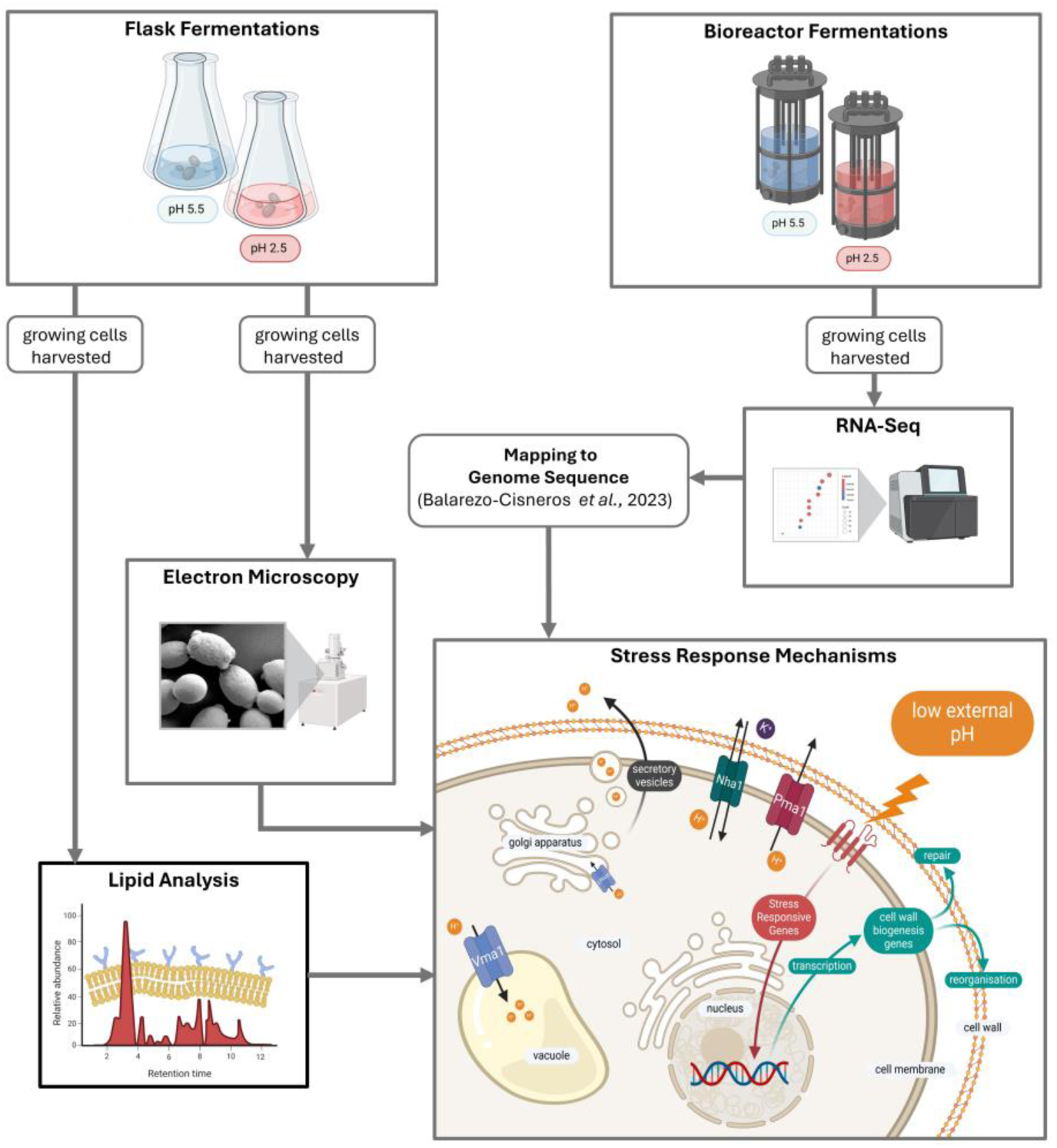
Experimental strategy of this study. To investigate the transcriptional and cellular response to low pH in *M. bulderi,* controlled bioreactor fermentations maintained at either pH 5.5 or 2.5. RNA-seq analysis was carried out to identify expressional changes that are related to the high growth of *M. bulderi* at low pH. Flask fermentations were carried out in parallel and cell wall structure of cells was analysed by electron microscopy and plasma membrane lipid composition was investigated through LC-MS. A diagram of yeast stress response mechanisms is included. Membrane bound ATPase pumps, the primary of which is Pma1, are the main mechanism for export of H+ ions from the cytoplasm. The vacuolar ATPase Vma1 which is located in the membranes of organelles allows the cell to store H+ in the vacuole and the golgi apparatus allowing excretion of H+ ions via the secretory pathway. Low external pH causes damage to the cell wall and triggers stress pathways including the cell wall integrity (CWI) and high osmolarity glycerol (HOG) pathways that are responsible for activation of genes involved in reorganisation and repair of the cell wall.

## 2 Results and Discussion

### 2.1 *M. bulderi* genome shows limited transcriptional reprograming under low pH conditions

To identify genes involved in the *M. bulderi* pH response, we investigated the gene expression profiles of three *M. bulderi* strains, namely CBS 8638, CBS 8639, and NRRL Y-27205, grown in bioreactors where the pH was maintained constant at either pH 5.5 or 2.5 throughout the fermentation. We first analysed the combined transcriptional response of all three strains to analyse the transcriptional changes at species level. In total, 500 genes exhibited strongly altered expression at a fold change (FC) greater than 2, p-adj<0.05 (*i.e.* 210 downregulated and 290 upregulated genes) in response to low pH. Additionally, a further 716 genes exhibited moderately altered expression at 1.5<FC<2, p-adj<0.05 (350 downregulated and 366 upregulated genes; p-adj<0.05) in response to low pH (Supplementary *data* 1). Expression of selected genes was validated by qPCR (Supplementary Fig. 1).

Enriched gene ontology (GO) terms showed that upregulated genes were predominantly linked to cell wall organization and biogenesis (Fig. 2A; and Supplementary Fig. 2), where both the CWI and HOG pathways play a major role. Conversely, among the downregulated genes, RNA processing, maturation, and ribosome biogenesis were enriched, suggesting a general slowdown of protein production (Fig. 2B; Supplementary Fig. 2). Three key genes in the CWI response *MID2, SLT2* and *RHO1,* responsible for signalling and regulation of the pathway, were all strongly upregulated in the *M. bulderi* species-wide response (Fig. 2C). Interestingly, the expression of *FKS1* and *FKS2,* two primary genes involved in cell wall repair and β-1,3-glucan synthesis that are upregulated in *S. cerevisiae* exposed to strong inorganic acids (de Lucena *et al.,* 2012) remained unchanged in *M. bulderi*. However, several cell wall biogenesis genes, such as the chitin synthase genes *CHS1*, *CHS3* and *CHS7*, were strongly upregulated in *M. bulderi*, as was *UTR2* which is involved in the transfer of chitin to β-glucans in the cell wall. Additionally, genes involved in remodelling of the β-glucan chains and protein mannosylation were strongly upregulated including *TOH1 NCW1*, *NCW*2, *KAR2*, *MNN2, KRE1, KRE6, KRE9, GAS1* and *GAS4*. In *S. cerevisiae, TOH1*, *GAS1*, *KRE6* have also been shown to be upregulated under exposure to strong inorganic acid (Kapteyn *et al.,* 2001; de Lucena *et al* 2012; de Lucena *et al*., 2015).

**Figure 2.**
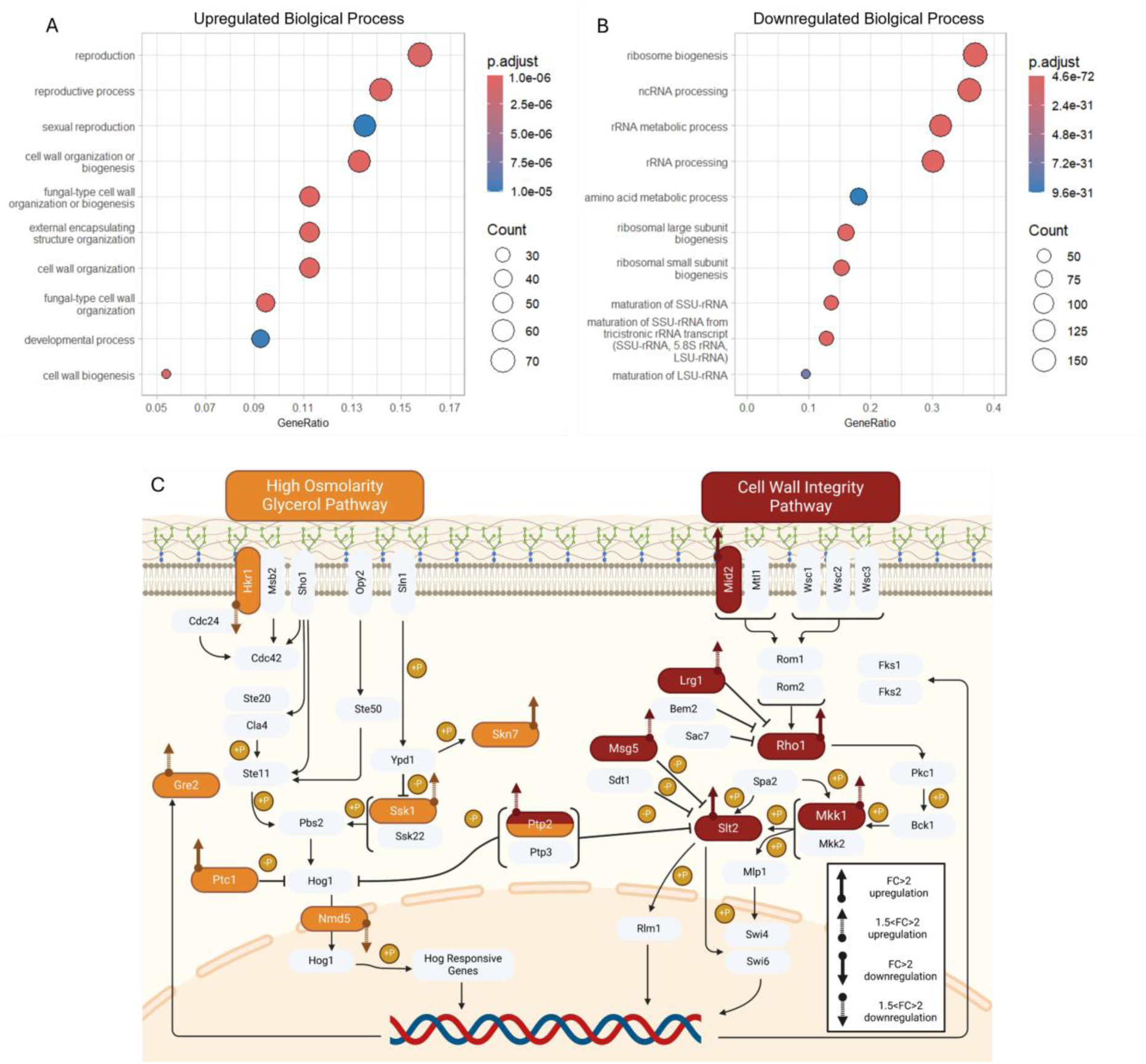
Enriched biological processes and differentially expressed genes involved in the CWI and HOG pathways. Plots showing Enriched GO biological process (BP) terms, in the combined *M. bulderi* species response at pH 2.5 vs pH 5.5, shown as upregulated genes (Panel A) and downregulated genes (Panel B). The gene ratio value is the ratio between the DE genes over the total of DE genes in the same category and the dot size corresponds to the number of DE genes linked to that function as a fraction of total DE. The GO terms reported are those significantly enriched in DE genes (p-adj<0.001). The DE genes at low pH involved in the high osmolarity glycerol (HOG) and cell wall integrity (CWI) pathways are highlighted in orange and red, respectively (Panel C).

The HOG signalling pathway is the primary response to osmotic stress in *S. cerevisiae* and several non-conventional yeast species (Van Wuystwinkel, *et al.,* 2000; Thomé, 2007; Ene *et al.,* 2015). It has been suggested that damage to the cell wall from acid stress can mimic osmotic stress through loss of cell structure (de Lucena *et al.,* 2012). In *M. bulderi* genes in the HOG pathway also showed dysregulation. *SKN7* a transcription factor that responds to osmotic stress (Fassler and West, 2011), was strongly upregulated at low pH, while *SSK1,* a signalling protein, and *GRE2*, a 3-methylbutanal reductase, were both moderately upregulated. *HKR1,* which codes for an osmosensing protein and activator of the HOG pathway, was moderately downregulated. In addition, *PTC1* an inhibitor of *HOG1* was strongly upregulated and *NMD5* a nuclear transporter involved in the nuclear localisation of Hog1p was moderately downregulated, which suggests an overall dampening of the HOG pathway. *CRZ1,* a transcription factor activated in the calcineurin pathway, a calcium-mediated signalling pathway that responds to environmental stress, was also moderately upregulated in *M. bulderi.* Gene targets of *CRZ1* were also upregulated. This included the moderate upregulation of cell wall related genes *SCW10*, *SCW11* and *KRE6* and strong upregulation of *KRE6* and *CRH1*. In *S. cerevisiae*, the CWI and HOG pathways overlap with the calcineurin pathway, when cells are exposed to low pH (Cyert, 2003; de Lucena *et al.,* 2012). It has been previously shown that *CRZ1* is important in the low pH response of yeasts. In *Candida glabrata CRZ1* deletants showed alteration in membrane lipid composition and lower viability under low pH stress (Yan *et al.,* 2016). In *S. cerevisiae CRZ1* is known to regulate cell wall synthesis genes *FKS1* and *FKS2* in response to cell wall damage (Cyert, 2003; del Lucena *et al* 2012), however, in *M. bulderi,* no upregulation of *FKS1* and *FKS2* was seen. It has been found previously that genes involved in the general stress response (GSR) are differentially regulated in *S. cerevisiae* under low pH stress (Kapteyn *et al*., 2001; De Melo *et al*., 2010; de Lucena *et al*., 2015). Few genes in the GSR were DE in *M. bulderi* and shared with *S. cerevisiae* response (4 out of 29 DE genes). These included *YGP1*, *SPI1, HSP150* and *GRE2,* all exhibiting a moderate increase in expression (Kapteyn *et al*., 2001; De Melo *et al*., 2010).

Among the enriched GO terms for upregulated genes, were some involved in sexual reproduction. Among them, and strongly upregulated, were the inducers of meiosis *IME1* and *IME2,* spore formation genes *SPO24* and *SPO74* and a-type mating hormone receptor *STE3.* It has been previously shown that *M. bulderi* has a very poor sporulation efficiency and only forms two globular ascospores (Middlehoven *et al.,* 2000). It is possible that low pH, being a stressor, triggers meiosis to produce tetrads. In our hands, however, we were unable to induce sporulation in this species (*i.e.* we could not see any tetrads or dyads).

GO enrichment analysis for cellular component showed a substantial portion of upregulated genes localized within the vacuole (Supplementary Fig. 2C). Specifically, genes involved in the transport across vacuolar membranes, namely *VVSI, SIT1*, *CTR2, AVT1, OPT2*, and *TPO4,* were all strongly upregulated at pH 2.5. In *S. cerevisiae, OPT2* is responsible for maintaining lipid asymmetry and maintenance of a charge gradient across membranes including the vacuolar membrane (Yamauchi *et al.,* 2015)*. VMA13,* a subunit of the V-ATPase required for its function, and *PMP2,* a regulator of membrane bound ATPases, were both strongly upregulated under low pH in *M. bulderi*. The V-ATPase has been shown to have a significant role in regulating both vacuolar and cytosolic pH in *S. cerevisiae* (Martínez-Muñoz & Kane, 2008). The upregulation of these genes suggests that the vacuole is a major player of the *M. bulderi* pH homeostasis response.

Among the cohort of *M. bulderi* species-specific genes, previously described in Balarezo-Cisneros et al., 2023 (*i.e*. those with no orthologs in any other species), 57 were DE, of which 34 strongly and 23 moderately. We mined our published dataset of AlphaFold structure and DeepFRI GO term predictions made for the *M. bulderi* (Balarezo-Cisneros *et al*., 2023), and found six upregulated genes (*KB390A01580*, *KB390A01870*, *KB390A02760*, *KB390A04940*, *KB390A05080* and *KB390A05760*), with an associated AlphaFold predicted functional annotation. Three genes were strongly upregulated including two predicted membrane transporters (*KB390A01870* and *KB390A05760*) and one signalling receptor binding protein (*KB390A05080*). The other three genes were moderately upregulated including two predicted membrane transporters (*KB390A01580* and *KB390A04940*) and one DNA binding protein (*KB390A02760*). These may be novel transporters that evolved specifically in the *Maudiozyma bulderi* lineage.

Overall, *M. bulderi* showed dysregulation of stress response pathways, including the CWI and HOG, similarly to *S. cerevisiae* in response to low pH. However, the overlap of response between these two species was somewhat limited, involving different genes. Additionally, unlike what has been reported in *S. cerevisiae*, we did not find many genes in *M. bulderi* with expressional changes associated with the GSR. This relatively limited transcriptional adjustment may reflect the fact that *M*. *bulderi* is better adapted to acid stress compared to *S. cerevisiae*.

### 2.2 The differential expression profile between *M. bulderi* strains recapitulates their phenotypic differences

We compared the transcription profiles of the three different *M. bulderi* strains, CBS 8638, CBS 8639 and NRRL Y-27205, at both pH 5.5 and 2.5. For all strains, the pH accounted for the majority of the variance in the RNA-seq data as shown by the PCA analysis (Fig. 3A). There was no strain effect between CBS 8638 and CBS 8639 strains, the PCA showed that CBS 8638 and CBS 8639 cluster together while NRRL Y-27205 shows a different profile at both pH 2.5 and 5.5 (Fig. 3A). Phenotypically, strains CBS 8638 and CBS 8639 exhibit more similar growth capabilities when exposed to strong inorganic acid, compared to NRRL Y-27205 profile (Balarezo-Cisneros, 2023). Overall, it is likely that CBS 8638 and CBS 8639 share a common regulatory response to stress conditions distinct from NRRL Y-27205. NRRL Y-27205 showed by far the greatest number of significantly DE genes (*i.e.* 1821), followed by CBS 8638 (*i.e.* 932) and CBS 8639 (*i.e.* 164; Supplementary Table 1). The results of GO enrichment analysis show important differences for each strain individually (Supplementary Fig. 3, 4 and 5). Differences in expression were also seen in many genes involved with pH homeostasis. Most notably a strong upregulation of *PMA1* was seen in NRRL Y-27205 at pH 2.5, while expression of this gene remained unchanged in CBS 8638 and CBS 8639. In *S. cerevisiae* Pma1p is the primary proton pump involved in pH homeostasis, however upregulation of *PMA1* is thought to be largely post translational although its activation is linked in particular to cytosolic acidification upon low pH stress (Carmelo *et al*., 1997; Martínez-Muñoz & Kane, 2008; Dechant *et al*., 2014; de Lucena *et al*., 2020). ATP channels and activity are crucial for an effective low pH response, as pumps rely on ATP for energy, with Pma1p activity estimated to consume approximately 15% of the ATP produced in metabolically active yeast (Lucena *et al*., 2020). It has been suggested that in other species such as *Pichia anomala*, induction of ATP production via respiration is a strategy to combat low pH stress by increasing H+ efflux through ATPases such as Pma1p (Fletcher *et al.,* 2015). However, this is unlikely to be the case in *M. bulderi* as none of the three strains possess complete mitochondrial DNA and are unable to respire (Balarezo-Cisneros *et al.,* 2023). NRRL Y-27205, unlike the other two strains, also showed upregulation of polysaccharide metabolism genes, *TPS2* and *UGP1* which are involved in trehalose synthesis. Trehalose is recognized as a reserve carbohydrate in yeast, and it has been shown that accumulation of trehalose an energy source occurs during starvation (Winderickx *et al*., 1996; Estruch, 2000).

**Figure 3.**
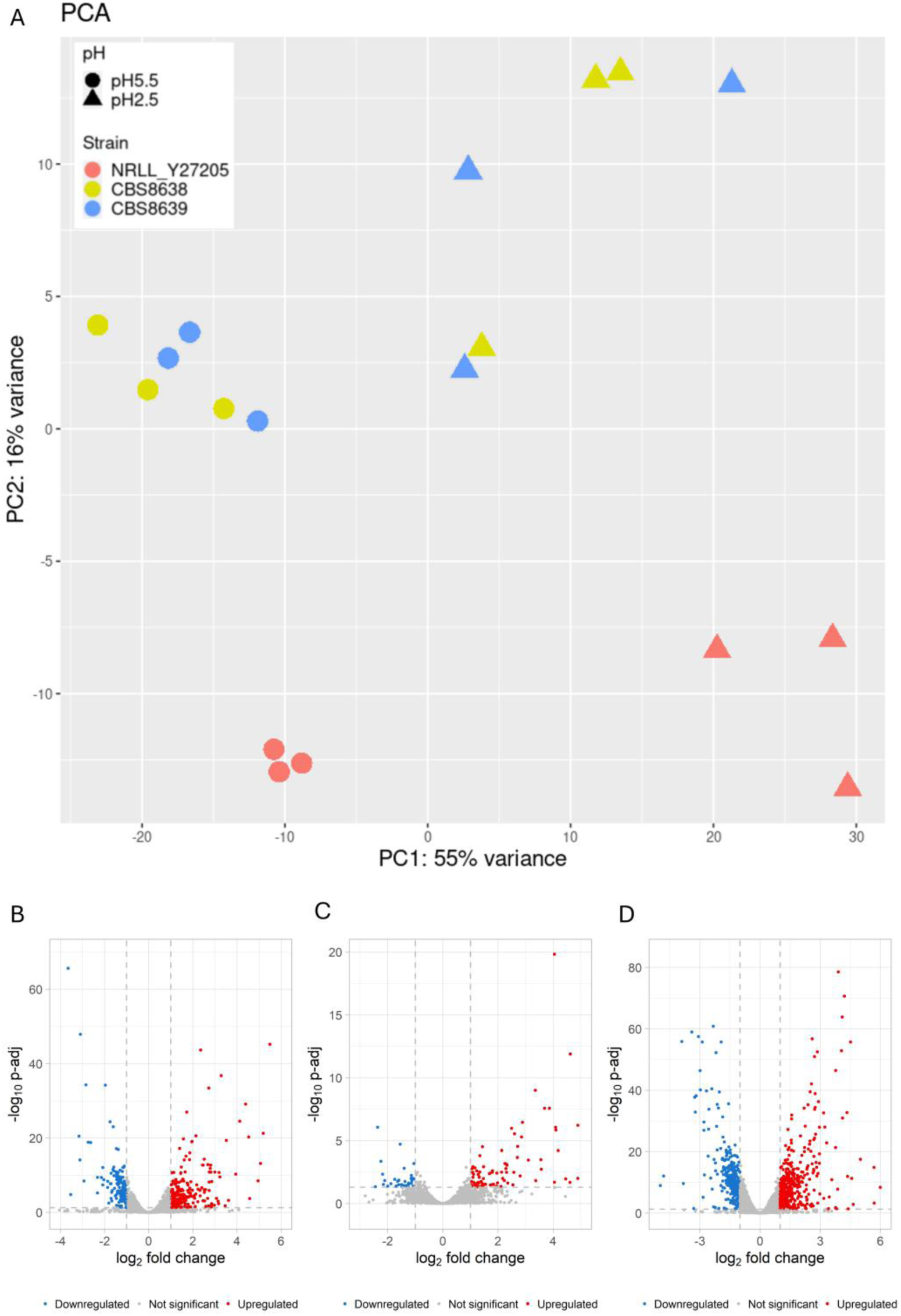
Expression profiles of *M. bulderi* strains. Principal Component Analysis plot of the gene expression profiles of *M. bulderi* strains (Panel A), and volcano plots of the RNA-seq analysis of *M bulderi* CBS 8638 (Panel B), CBS 8639 (Panel C) and NRRL Y-27205 (Panel D) grown on pH 5.5 vs 2.5. The highlighted genes in blue and red have FC >1.5 or FC<-1.5, with p-adj<0.05.

In NRRL Y-27205 at low pH, we saw greater activation of stress response pathways. For example, the DE genes in the CWI pathway included *MSG5* and *WSC2,* neither of which were DE in CBS 8638 nor CBS 8639. *PTP2* was also strongly upregulated in NRRL Y-27205. This gene is only moderately upregulated in CBS 8638 and not DE in CBS 8639. *WSC2* is an important sensor of damaged cell wall and low pH in the CWI pathway. *MSG5* and *PTP2* are involved in the suppression of *SLT2*, a transcription factor in the CWI, which was upregulated in both NRRL Y-27205 and CBS 8638 which suggests that negative feedback to reverse *SLT2* activation has been triggered. A similar picture was seen with genes responsive to activation of the HOG pathway, where NRRL Y-27205 shows upregulation of *GRE2, SSK1, OPY2, SIL1* and *TUP1,* none of which are DE in CBS 8638 or CBS 3639. This suggests that NRRL Y-27205 may be incurring damage to the cell structure, mimicking osmotic stress, to a greater extent than CBS 8638 and CBS 8639 (de Lucena *et al.,* 2012). Furthermore, in NRRL Y-27205 we observed a higher number of DE genes involved in the GSR at low pH (*GCY1*, *HSP42*, *HSP150, GRE2, YGP1* and *SPI1*) when compared to those in CBS 8638 (*HSP150, SPI1* and *YGP1)* and CBS 8639 (none).

Different combinations of heat shock proteins were strongly upregulated in the three strains at low pH. *HSP26* was DE in all three strains, *HSP150* was DE in CBS 8638 and NRRL Y-27205 while *HSP42, HSP78* and *HSP104* were DE only in NRRL Y-27205. Previous studies have shown that *HSP26* and *HSP150* are upregulated in *S. cerevisiae* in response to low pH (Carmelo and Sá-Correia, 1997; Kapteyn *et al*., 2001) and *HSP104* has been shown to be involved in the cell response to various stresses but it’s role in the acid response has not been characterised (Sanchez *et al.,* 1992; Amorós and Estruch, 2001). In NRRL Y-27205 we also saw a strong upregulation of *HXT13* (glucose transporter) and *SKS1* (regulation of glucose transport). Previous studies showed that low external pH can mimic conditions of glucose starvation in *S. cerevisiae*, and that genes related to glucose transporters and adaptation to low glucose concentrations were upregulated (De Melo *et al*., 2010).

Overall, NRRL Y-27205’s response to low pH showed much greater transcriptional disruption and included activation of several genes associated with pH stress that has been also described in *S. cerevisiae* which were not observed in CBS 8638 or CBS 8639. Combined with the phenotypic data at low pH, it is clear that CBS 8638 and CBS 8639 are better adapted to grow efficiently at low pH, while NRRL Y-27205, as well as *S. cerevisiae*, faces more pronounced challenges at low pH, resulting in a more widespread response compared to CBS 8638 and CBS 8639.

### 2.3 Structural variation contributes to expressional differences among strains

The telomere-to -telomere assembly of the *M. bulderi* genome showed that CBS 8638 and CBS 8639 possess fully collinear genomes, while NRRL Y-27205 has an inversion in chromosome VII (Balarezo-Cisneros *et al*., 2023). Three out four genes flanking the breakpoint, *AQY1* (a aquaporin water channel), *CYT2* (involved in the maturation of cytochrome c1) and *YKL162C* (an uncharacterised protein that localises to the mitochondrion) are common to all three strains. NRRL Y-27205 has one different gene at the breakpoint, namely *NR270G02130*, to the other two strains, which both possess *KB390G02120* (Suplementary Fig., 6A). *NR270G02130* is present in CBS 8638 and CBS 8639 but elsewhere on chromosome VII. Chromosomal rearrangements can alter on the expression of local genes surrounding the breakpoints (Puig *et al.,* 2000; Naseeb & Delneri, 2012; Naseeb *et al.,* 2016) and such transcriptional differences can impact on the phenotype. Here, we checked if genes situated at the breakpoints of the chromosome VII inversion displayed DE in the three strains at pH 2.5 vs pH 5.5. We observed that in CBS 8638 and CBS 8639, none of the genes located at the breakpoints were significantly DE at low pH. However, in NRRL Y-27205, *AQY1, NR270G02130* and *CYT2* were significantly DE. *CYT2* was moderately upregulated and *AQY1* and *NR270G02130* were strongly upregulated in NRRL Y-27205 (Fig. 4). The molecular function of *NR270G02130* is unknown as is its contribution to pH stress. However, *AQY1* is known to be induced during spore formation in *S. cerevisiae*. When *AQY1* is deleted in *S. cerevisiae* spore viability is decreased but vegetative cell viability under osmotic stress conditions is increased. (Bonhivers *et al.,* 1998; Sidoux-Walter *et al.,* 2004). The increased expression of *AQY1* in NRRL Y-27205 could therefore play a part in its reduced pH tolerance compared to CBS 8638 and CBS 8639, possibly by increasing the permeability of the cells via the formation of water channels. Alternatively, this may suggest that sporulation is induced in NRRL Y-27205 by low pH. The impact of *CYT2* upregulation is not clear in *M. bulderi* as none of the strains are able to respire.

**Figure 4.**
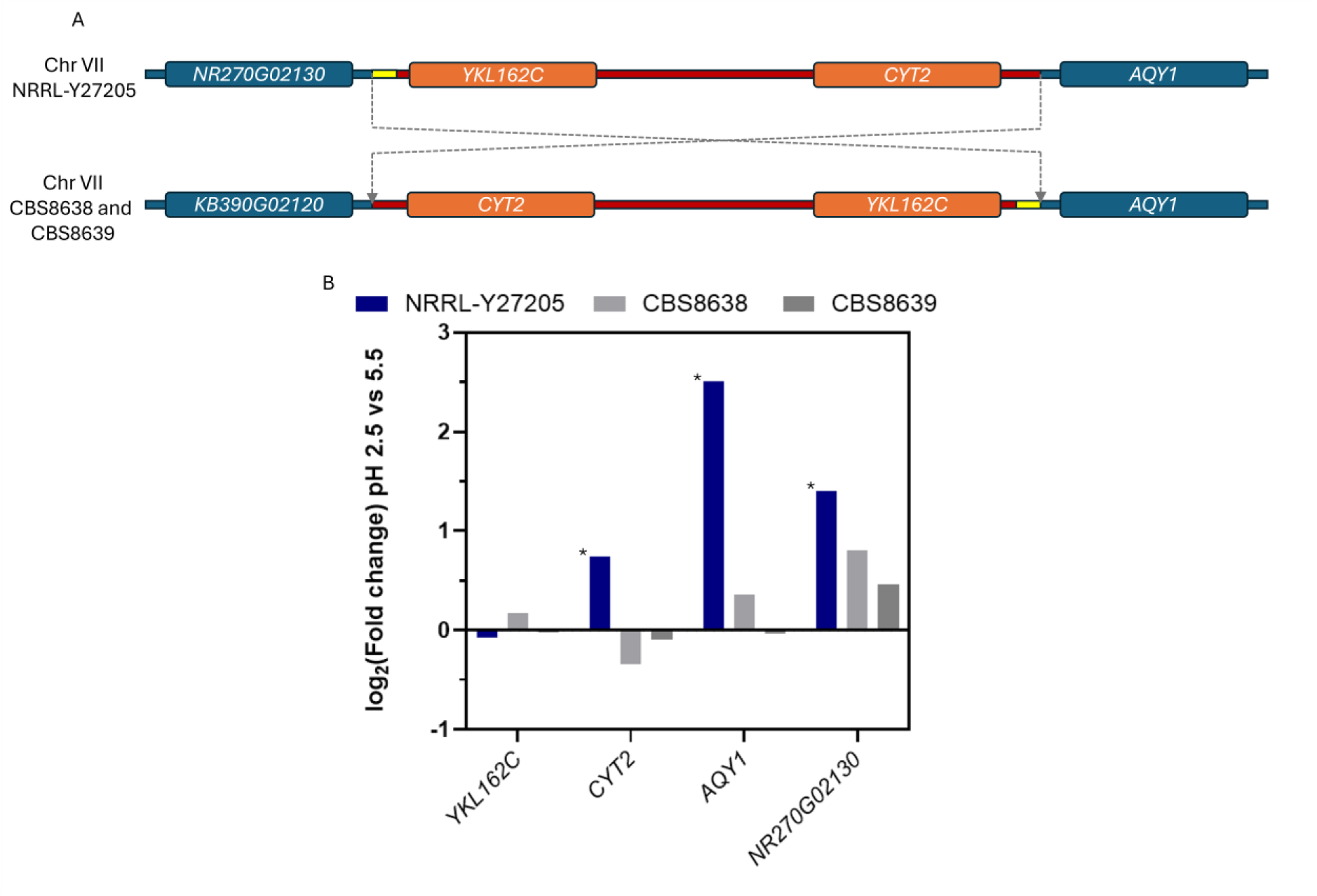
Expressional changes of genes neighbouring the inversion breakpoint. Diagram representing the inversion in chromosome VII found in *M. bulderi* strain NRRL-Y27205 compared to CBS8638 and CBS8639 with the flanking genes highlighted (Panel A). Chart showing the relative log2(FC) of each gene flanking the breakpoint in each strain at pH 2.5 vs pH 5.5 (Panel B). Bars marked with an asterisk denote significant differential expression (padj<0.05).

### 2.4 The *M. bulderi* cell wall shows limited physical disruption under low pH

The *M. bulderi* transcriptome revealed the majority of DE genes were associated with cell wall synthesis and restructuring. To investigate the effect of low pH on cell wall structure, *M. bulderi* cells exposed to pH 5.5 and pH 2.5 were imaged by scanning electron microscopy, alongside with *S. cerevisiae* industrial strain NCYC 505 cells as a control. *S. cerevisiae* cells imaged in permissive pH of 5.5 showed minimal signs of cell wall stress, however, many cells grown at pH 2.5 expressed a severely distorted morphology with large cell wall invaginations (Fig. 5 A and B; Supplementary Fig. 6 A and B). These results are consistent with previous literature showing similar damage to *S. cerevisiae* cell wall structure at low pH in both strong inorganic and weak organic acid (de Lucena *et al*., 2015; Dong *et al*., 2017). Several *S. cerevisiae* cells also showed globular cell surface structures at low pH, similar to those observed as a result of oxidiative stress in *Y. lypolitica* (Arinbasarova *et al.,* 2018) or as results of antifungal drug stress on *Candida albicans* (Rukayadi and Hwang, 2007). On the other hand, all *M. bulderi* strains showed no apparent damage at pH 5.5, and only a few CBS 8638 and NRRL Y-27205 cells were covered in globular structures at pH 2.5 (Fig. 5 C, D, G and H, Supplementary Fig. 6, C, D, G and H). Cells of CBS 8639 showed no apparent signs of cell wall damage at both pH 5.5 and pH 2.5 (Fig. 5, E and F; Supplementary Fig. 6 E and F). Studies in *S. cerevisiae* have linked globular structures appearing on the cell surface during protoplast regeneration with the secretion of Ygp1p and Hsp150p, two glycoproteins that attach to the cell wall (Pardo *et al.,* 1999) and are known to be upregulated in response to osmotic stress (Zueco, 2001). In our dataset, both *YGP1* and *HSP150* were upregulated at low pH in NRRL Y-27205 and CBS 8638 but not in CBS 8639, which showed the least cell wall damage. It is possible that the overexpression of these genes contributes to the globular structures seen on *M. bulderi* cell surface.

**Figure 5.**
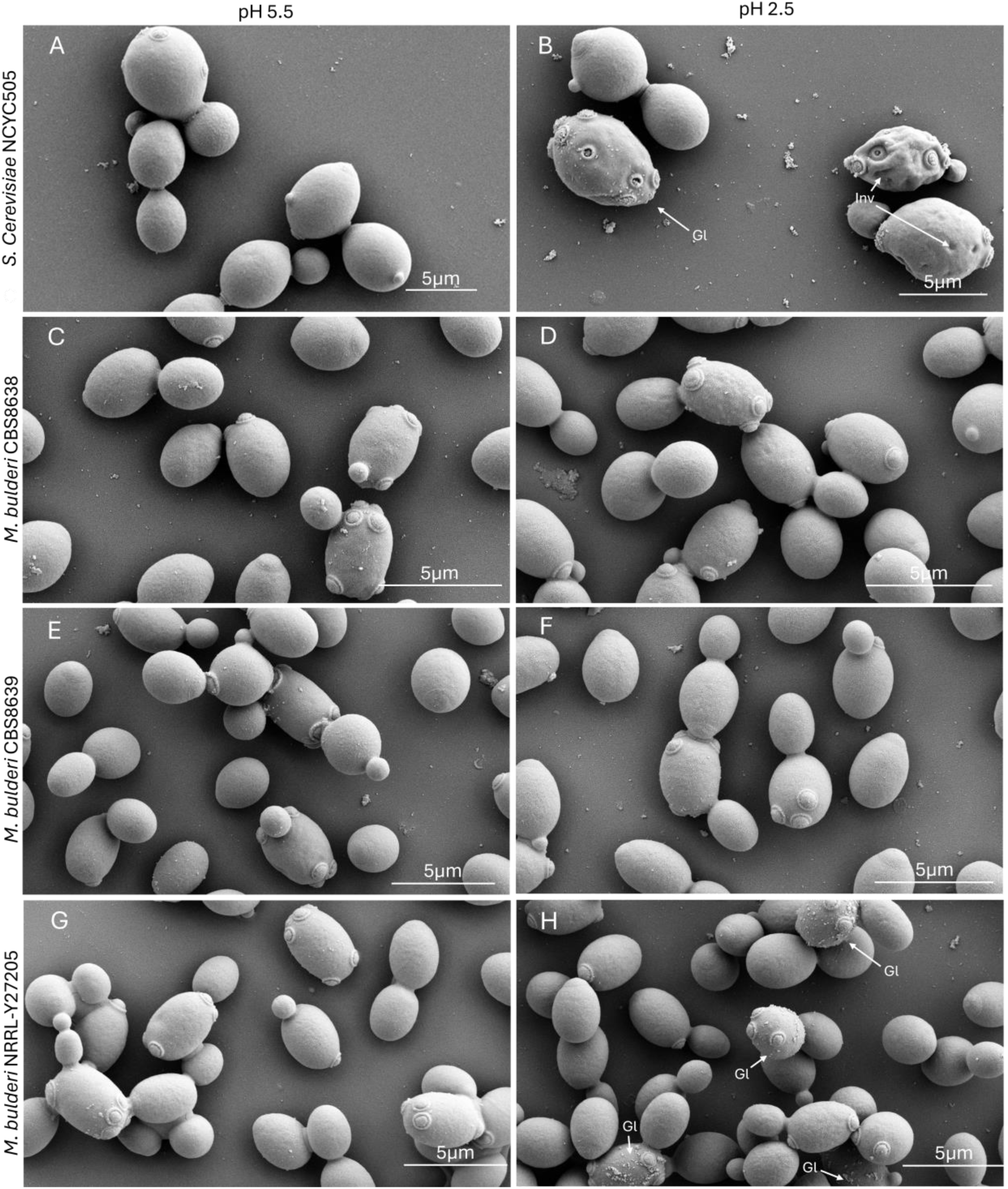
Cell wall morphology of *M. bulderi* and *S. cerevisiae* under low pH. Scanning electron micrographs of *Saccharomcyes cerevisiae and Maudiozyma bulderi* grown in minimal media under the following conditions: Panel A: NCYC505 pH 5.5, Panel B: NCYC505 pH 2.5, Panel C: CBS8638 pH5.5, Panel D: CBS8638 pH 2.5, Panel E CBS8639 pH 5.5, Panel F: CBS8639 pH 2.5, Panel G: NRRL Y-27205 pH 5.5, Panel H: NRRL Y-27205 pH 2.5. Damage to the cell wall including globular structures (Gl) and invaginations (Inv) are marked with arrows.

Overall, all the *M. bulderi* strains showed much less disruption to the cell wall compared to *S. cerevisiae* at low pH, and these findings are consistent with the growth phenotypes observed at low pH for these stains. Among the *M. bulderi* strains, NRRL Y-27205 appeared to have the most affected cell wall morphology which is consistent with it showing the poorest growth at low pH (Balarezo-Cisneros *et al*., 2023).

### 2.5 In *M. bulderi* CBS 8639 the membrane lipid composition is stable at low pH

The lipid composition is a trade-off for multiple stress factors ranging from temperature tolerance to membrane fluidity and organic acids permeability that can cross the membrane. Ergosterol has a particularly important role impacting membrane permeability and facilitating the localisation of membrane bound proteins (Suchodolski *et al*., 2019). To investigate any differences in the lipid composition of *M. bulderi* when grown at permissive pH (5.5) vs low pH (2.5), dried cells of *M. bulderi* CBS 8639 was compared with *S. cerevisiae,* strain NCYC 505, harvested during mid-log growth phase under both pH conditions. CBS 8639 cells showed a significantly lower mass of lipid as a fraction of dried cell weight when compared with *S. cerevisiae* at both pH 5.5 and 2.5 (Supplementary Fig. 7). Both species showed no significant change to total lipid mass as a proportion of dried cell weight between pH 2.5 and pH 5.5.

*S. cerevisiae* showed a significant decrease in glycerolipids from 64.8% to 43.8% (Multiple T-test p-adj <0.05). Ergosterol also increased, although the difference remained below the significant threshold (p-adj <0.05). In *S. cerevisiae g*lycerolipids such as triacylglycerol are used as energy storage in yeast cells (Illingworth *et al.,* 1973; Sorger and Daum, 2003). Additionally, H+ efflux via ATPases accounts for the majority of energy use under low pH stress (Ullah *et al.,* 2013). The reduction of glycerolipids seen in *S. cerevisiae* likely represents a consumption of storage energy to power pH homeostasis. Ergosterol has been shown to impact acid tolerance in yeasts. *MED3* deletant mutans of *C. glabrata* that were unable to synthesise ergosterol showed reduced viability in response to strong organic acid (Lin *et al.,* 2017). *S. cerevisae* has also been shown to better tolerate organic acids such as lactic acid with increased levels of ergosterol (Ferraz *et al.,* 2023). Interestingly, *M. bulderi* CBS 8639 showed no significant change in composition of lipids between ph 5.5 and 2.5 (Fig. 6), suggesting that *M. bulderi* membrane is stable at this pH and does not require lipid restructuring. Limited transcriptional change was observed in *M. bulderi* CBS 8639 related to lipid synthesis. However, *SFK1,* a membrane bound phospholipid flippase, responsible was strongly upregulated at low ph. It has been shown in *S. cerevisiae* that *SFK1* is upregulated in response to low pH and is required to maintain membrane impermeability and the retention of ergosterol in membranes (Mioka *et al.,* 2018; Kishimoto *et al.,* 2021). It is therefore likely that *M. bulderi* is already well adapted to grow at the pH tested in this study and does not rely on alterations to lipid structure to maintain growth at low pH.

**Figure 6.**
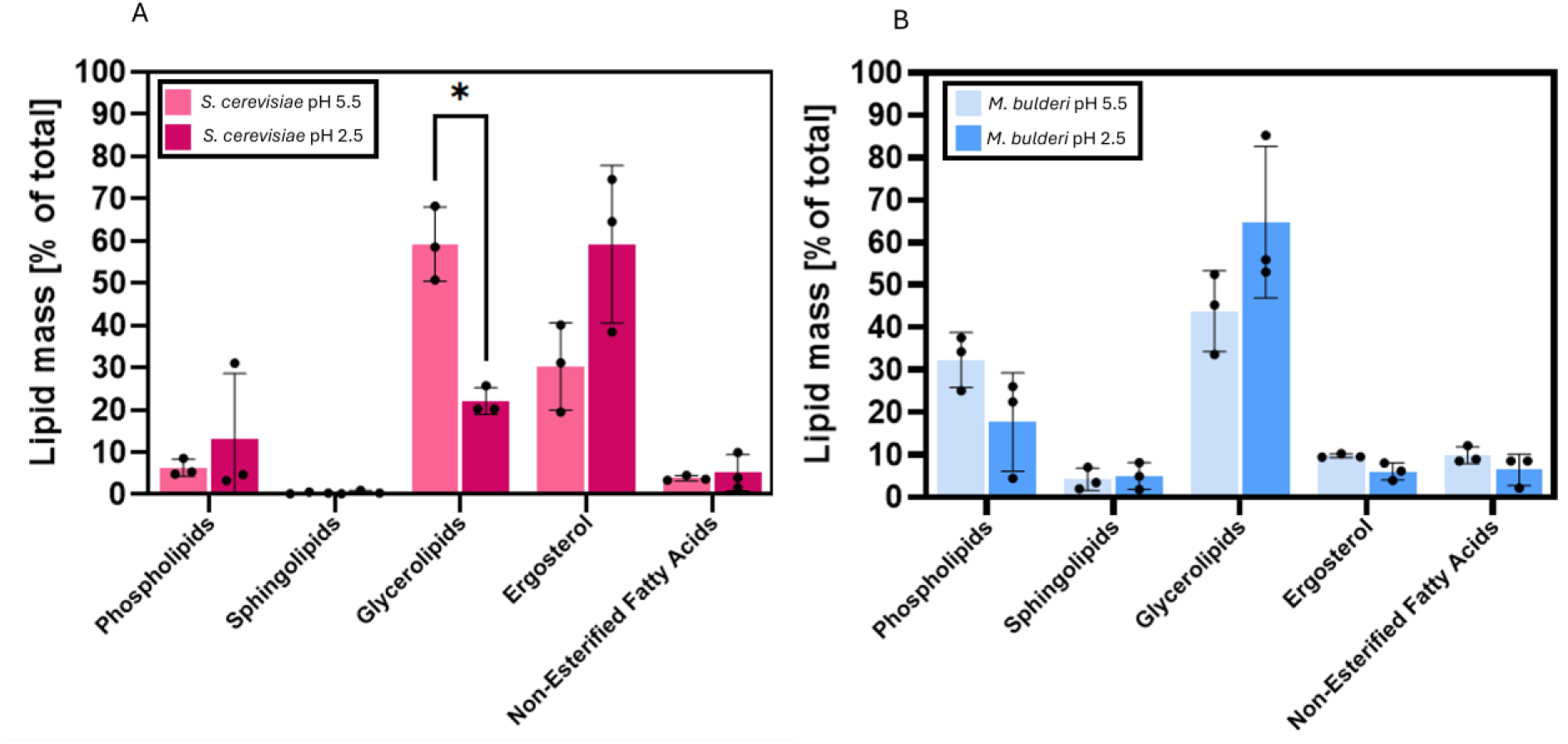
Lipidomics profiling of *M. bulderi* and *S. cerevisiae*. Lipid mass expressed as % of total lipids in *S. cerevisiae* NCYC505 grown at permissive pH 5.5 and acid pH 2.5 (Panel A) and lipid mass expressed as % of total lipids in *M. bulderi* CBS8639 grown at permissive pH 5.5 and acid pH 2.5 (Panel B).

Moreover, we identified a subset of differentially expressed (DE) genes which are unique to *M. bulderi* (*i.e.* lacking homologs in other species), that may play crucial roles in the species’ survival under stress. Comparative analysis between strains revealed distinct transcriptional responses, recapitulating phenotypic diversity among *M. bulderi* populations. Additionally, we showed that, unlike *S. cerevisiae*, *M. bulderi* cell wall is not damaged by low pH and the lipid composition also remains unchanged.

Overall, our study sheds light on the pathways underpinning the remarkable tolerance of *M. bulderi* to acidic environments. These findings not only expand our understanding of microbial adaptation to extreme conditions but also underscore the importance of genetic and transcriptional variability in shaping the stress responses in yeast species.

## 3 Conclusions

Our investigation revealed that under low pH, *M. bulderi* upregulated genes associated mainly with cell wall restructuring and synthesis. The upregulated genes showed some similarities in the gene expression landscape of those described in *S. cerevisiae,* however, more limited dysregulation was observed in *M. bulderi,* with reduced activation of the CWI and HOG pathways. The individual strains of *M. bulderi* displayed distinct transcriptional responses that correlated with the differences in phenotype observed in the strains at low pH. CBS 8638 and CBS 8639 exhibited a more closely shared response, with relative transcriptional stability, while NRRL Y-27205 exhibited the strongest dysregulation and also the weaker ability to grow at low pH (Balerzo-Cisneros, dt al, 2023). We also showed that the chromosomal inversion present in NRRL Y-27205 is affecting the expression of the genes located at chromosomal breakpoints, being upregulated compared to those in CBS 8638 and CBS 8639, suggesting a phenotypic impact. Moreover, we identified a subset of differentially expressed (DE) genes which are unique to *M. bulderi* (*i.e.* lacking homologs in other species), that may play crucial roles in the species’ survival under stress. By coupling the transcriptional data with protein functional predictions, this study identifies potential novel membrane transporters, specific to *M. bulderi*, with a role in low pH tolerance. Our imaging data also showed that at low pH *M. bulderi* has greater structural stability with a cell wall that is more resistant to damage than *S. cerevisiae.* Additionally, the membrane lipid composition of CBS 8639 was largely unchanged under low pH unlike *S. cerevisiae.* Overall, this research sheds light on the cellular pathways involved in the acidophilic adaptation of *M. bulderi*, it shows that the strain best adapted to low pH has transcriptional stability to changing pHs, and highlights the potential importance of structural variants in shaping adaptive responses. These findings not only expand our understanding of microbial adaptation to extreme conditions but also underscore the importance of genetic and transcriptional variability in shaping the stress responses in yeast species.

## 4 Materials and Methods

### 4.1 Strains and media

Three *Maudiozyma bulderi* strains used in this study CBS 8638, CBS 8639, and NRRL Y-27205 were obtained from the Agricultural Research Service (ARS-NRRL) culture collection. *Saccharomyces cerevisiae* NCYC 505 was obtained from the National Collection of Yeast Cultures (NCYC). The yeasts were cultured and maintained on YPD agar medium (15 g/L agar, 10 g/L yeast extract, 20 g/L peptone, 20 g/L glucose) or in SD minimal media (Yeast nitrogen base with ammonium sulphate and amino acids 6.7g/L)

### 4.2 Bioreactor fermentation for RNA extraction

Bioreactor fermentations for RNA extraction were conducted using one litre stirred tank bioreactors (Multifors, Infors-HT, Bottmingen, Switzerland). All fermentation experiments were carried out with cells grown in 500 mL of SD minimal media supplemented with 20g/l glucose. Fermentations were inoculated with cells from YPD overnight culture, washed twice in sterile deionised water, to achieve an initial optical density (OD) of 0.1. Three replicates of each strain were then cultured in 500 mL of YNB supplemented with 20g/l glucose and adjusted to either pH 5.5 or 2.5 with H_2_SO_4_. The pH was maintained at 5.5 and 2.5 using either 10 M NaOH or 1 M H_3_PO_4_ for pH adjustment. Temperature was maintained at 25°C, with agitation set at 300 rpm. For RNA extraction, 10-20 mL of cells with OD between 0.7 and 1.0 were harvested and promptly snap-frozen with liquid nitrogen.

### 4.3 Total RNA Extraction and Quantitative RT-PCR

Total RNA was extracted from 1×10^7 cells of three biological replicates harvested at the logarithmic growth phase (OD600 0.5-1.0) using the RNeasy Mini Kit (QIAGEN, Germany) following the manufacturer’s protocol for enzymatic digestion of the cell wall followed by lysis of spheroplasts. The concentration of lyticase was optimized to 50U per 10^7^ cells for efficient cell lysis. To eliminate genomic DNA contamination, an additional DNase treatment was performed using the RNase-free DNase set (QIAGEN, Germany) according to the manufacturer’s instructions. The extracted RNA was quantified using a NanoDrop Lite Spectrophotometer (Thermo Fisher Scientific, United States). One microgram of total RNA was reverse-transcribed into cDNA using the QuantiTect Reverse Transcription Kit (QIAGEN, Germany) following the manufacturer’s protocol. Optimized qPCR reactions consisted of 2 ng of cDNA, 3 pmol of each primer, and 5 µl of iTaq Universal SYBR Green Supermix 2X in a final volume of 10 µl. Reactions were run on a Roche LightCycler real-time System for 35 cycles with the following conditions: 15 s at 95 °C, 30 s at 55 °C, and 30 s at 72 °C

### 4.4 Illumina HiSeq library preparation and sequencing

Libraries were prepared from total RNA using the TruSeq Stranded mRNA Library Prep Kit (Illumina, Inc.) following the manufacturer’s instructions. Sequencing was conducted on an Illumina HiSeq 4000 instrument at the Genomic Technologies Core Facility, University of Manchester. Quality control of the reads was performed using FastQC (v0.11.9) (https://www.bioinformatics.babraham.ac.uk/projects/fastqc/).

Genes were aligned to the *M. bulderi* CBS 8639 annotated genome (Balarezo-Cisneros *et al*., 2023) using the STAR aligner version 2.5.3a (Dobin *et al*., 2013) and CBS 8639 served as the standard assembly for all three strains. Differential gene expression analysis was conducted based on the negative binomial distribution using DESeq2 (Love *et al*., 2014) in R version 4.3.2. Volcano plots were generated with DESeq2 output for each strain in R using the ggplot2 package. Genes exhibiting a statistically significant difference in expression, with a p-adjusted value below 0.05 and a fold change greater than 1.5 or 2, were selected for further analysis.

### 4.5 Functional enrichment analysis

Enrichment analysis was performed using the R package clusterProfiler (Yu *et al*., 2012) which included Gene Ontology (GO) terms and Kyoto Encyclopedia of Genes and Genomes (KEGG) pathways (Kanehisa & Goto, 2000). GO terms with FC>1.5 or -1.5>FC and adjusted p-values (Padj) < 0.05 were deemed significantly enriched. The GO analysis encompassed three categories: Biological Process (BP), Molecular Function (MF), and Cellular Component (CC). Differentially expressed (DE) genes were categorised as up or down-regulated and enrichment analysis was conducted. Only genes that had been functionally annotated with *S. cerevisiae* homologs were considered for this analysis. GO enrichment analysis was carried out on the combined *M. bulderi* response and on each of the three strains separately.

### 4.6 Electron microscopy

Shake flask cultures of *M. bulderi* strains CBS 8638, CBS 8639 and NRRL Y-27205 and *S. cerevisiae* NCYC 505 of 50ml of SD minimal media supplemented with 20g.l-1 of glucose and adjusted to either pH 5.5 of pH 2.5 with H_2_SO_4_ with three replicates for each strain. Flasks were inoculated with a single colony and incubated at 25C with shaking of 180rpm for approximately 12 hours. Cells were harvested in mid-log growth phase at an OD_600_ of between 0.5-0.7. Cell suspensions were settled on glass coverslips coated with L-polylysine for 30 min, then a droplet (1/10 of suspension volume) of 25% glutaraldehyde stock solution was added to each coverslip directly into cell suspension volume for chemical fixation over 1h at room temperature. After that coverslips were washed in 0.1M Cacodylate buffer (pH 7.2) to remove excess cells. Samples were fixed with 1% OsO4 in 0.1 Cacodylate buffer (pH 7.2), washed with distilled water and incubated with 1% uranyl acetate in water for at least 1h. After washing with distilled water, coverslips were brought through the ascending concentrations of ethanol until 100%. Then the ethanol was exchanged for HMDS by the incubation with HMDS/ethanol mixes with ascending HMDS concentration (up to 100%). Finally, coverslips were air dried. Dry samples were sputter coated in Leica ACE600 with Au/Pd layer of 16nm thickness. Cells were examined using an Apreo 2S FEG SEM (Thermo Fisher Scientific, United States) at 5kV acceleration voltage in Optiplan lens mode.

### 4.7 Lipid analysis

Lipid characterisation was carried out on dried cells harvested from *M. bulderi* strain CBS 8639 and *S. cerevisiae* NCYC 505. Shake flask fermentations were carried out with the same method as described above for electron microscopy, with the only difference of using 200ml of culture for each shake flask. Following harvest, cells were pelleted by centrifugation and washed with sterilized deionised water twice and dried in an oven at 60°C for 24hrs. Dried cells were sent to Creative Proteomics (New York, USA) where the samples were analysed using the following protocol: yeast samples on dry ice were spiked with 20 microliters of an internal standard and calibration mixture consisting of 500 picograms/microliter of dimyristoyl phospholipids (PG, PE, PS, PA,PC), SM(35:1), Cer(30:1), Cholesteryl ester(14:0) and TG(45:0). The spiked samples were combined with 300 μL of -20C chilled 75% methanol containing 1 mM BHT and homogenized using ceramic beads. To each homogenate, one mL of MTBE was added, and samples were then vortexed for 60 minutes at room temperature. 230 microliters of water was added, and the samples were further vortexed for an additional 10 minutes. After centrifugation for 10 minutes, the supernatants were collected to new test tubes and precipitated proteins were re-extracted as above after the addition of 250 ul of water. Pooled extracts were combined, dried overnight in a speedvac, and resuspended in 500 microliters of isopropanol. Full scan MS spectra at 100,000 resolution (defined at m/z 400) were collected on a ThermoScientific LTQ-Orbitrap XL mass spectrometer (ThermoFisher Scientific, Massachusetts, USA) in both positive and negative ionization modes. Scans were collected from m/z 200 to m/z 1200. For each analysis, 10 μL of sample was directly introduced by flow injection (no LC column) at 10 μL/min using an electrospray ionizaton source equipped with a heated ESI needle. A Dionex Ultimate 3000 HPLC (ThermoFisher Scientific, Massachusetts, USA) in served as the sample delivery unit. The sample and injection solvent were 2:1 (v: v) isopropanol: methanol containing 20 mM ammonium formate. The spray voltage was 4.5 kV, ion transfer tube temperature was 275 °C, the S-lens value was 50 percent, and the ion trap fill time was 100 ms Samples were analyzed in random order, interspersed by solvent blank injections. Following MS data acquisition, offline mass recalibration was performed with the “Recalibrate Offline” tool in Thermo Xcalibur software according to the vendor’s instructions, using the theoretical computed masses for the internal calibration standards. MS/MS confirmation and structural analysis of lipid species identified by database searching were performed using CID-MS/MS at 60,000 resolution and a normalized collision energy of 25 for positive ion mode, and 60 for negative ion mode. MS/MS scans were triggered by inclusion lists generated separately for positive and negative ionization modes. Lipids were identified using the Lipid Mass Spectrum Analysis (LIMSA) v.1.0 software linear fit algorithm, in conjunction with an in-house database of hypothetical lipid compounds, for automated peak finding and correction of 13C isotope effects. Peak areas of found peaks were quantified by normalization against an internal standard of a similar lipid class. The top ∼300 most abundant peaks in both positive and negative ionization mode were then selected for MS/MS inclusion lists and imported into Xcalibur software for structural analysis on a pooled sample as described above.

## Acknowledgments

The authors wish to thank the Genomic Technologies and Bioinformatic Core Facility and the Electron Microscopy Core Facility at the University of Manchester. The authors also wish to thank Federico Visinoni for his invaluable support in setting up the fermenters and for sharing his expertise on the fermentation processes. This work was supported by BBSRC-link grant (BB/T002123/1) awarded to D.D. with economic support provided by BP. A.H. is supported by a studentship from the Future Biomanufacturing Hub, funded by the Engineering and Physical Sciences Research Council (EPSRC) and Biotechnology and Biological Sciences Research Council (BBSRC), as part of UK Research and Innovation (grant EP/S01778X/1) and BP.

## Data Availability

The transcriptome data are deposited in ArrayExpress with the ID number E-MTAB-15136.

