## Supplementary Files for "Unveiling the genetic basis of the low pH response in the acidophilic yeast *Maudiozyma bulderi* as a potential host for biorefinery"

**Supplementary Table 1.** Number of DE genes individually in CBS 8638, CBS 8639 and NRRL Y-27205 strains above fold change 2 and between fold change 1.5 and 2.

**Strongly differentially expressed genes x>FC2**

| | Downregulated | | Upregulated | | total |
|---|---|---|---|---|---|
| | Annotated | *M. bulderi* specific | Annotated | *M. bulderi Specific* | DE |
| CBS8638 | 174 | 1 | 249 | 11 | 435 |
| CBS8639 | 34 | 2 | 87 | 8 | 131 |
| NRRL-Y27205 | 306 | 23 | 420 | 16 | 765 |

**Moderately differentially expressed genes FC1.5<x<FC2**

| | Downregulated | | Upregulated | | total |
|---|---|---|---|---|---|
| | Annotated | *M. bulderi* specific | Annotated | *M. bulderi Specific* | DE |
| CBS8638 | 266 | 2 | 222 | 7 | 497 |
| CBS8639 | 18 | 0 | 14 | 1 | 33 |
| NRRL-Y27205 | 531 | 23 | 496 | 6 | 1056 |

**Supplementary Figure 1. Validation of RNAseq on selected candidates**

Chart comparing RNAseq and qPCR data showing fold change in genes *AQY1* and NR2702130 in NRRL Y-27205 and CBS8638 between pH 2.5 and pH 5.5.

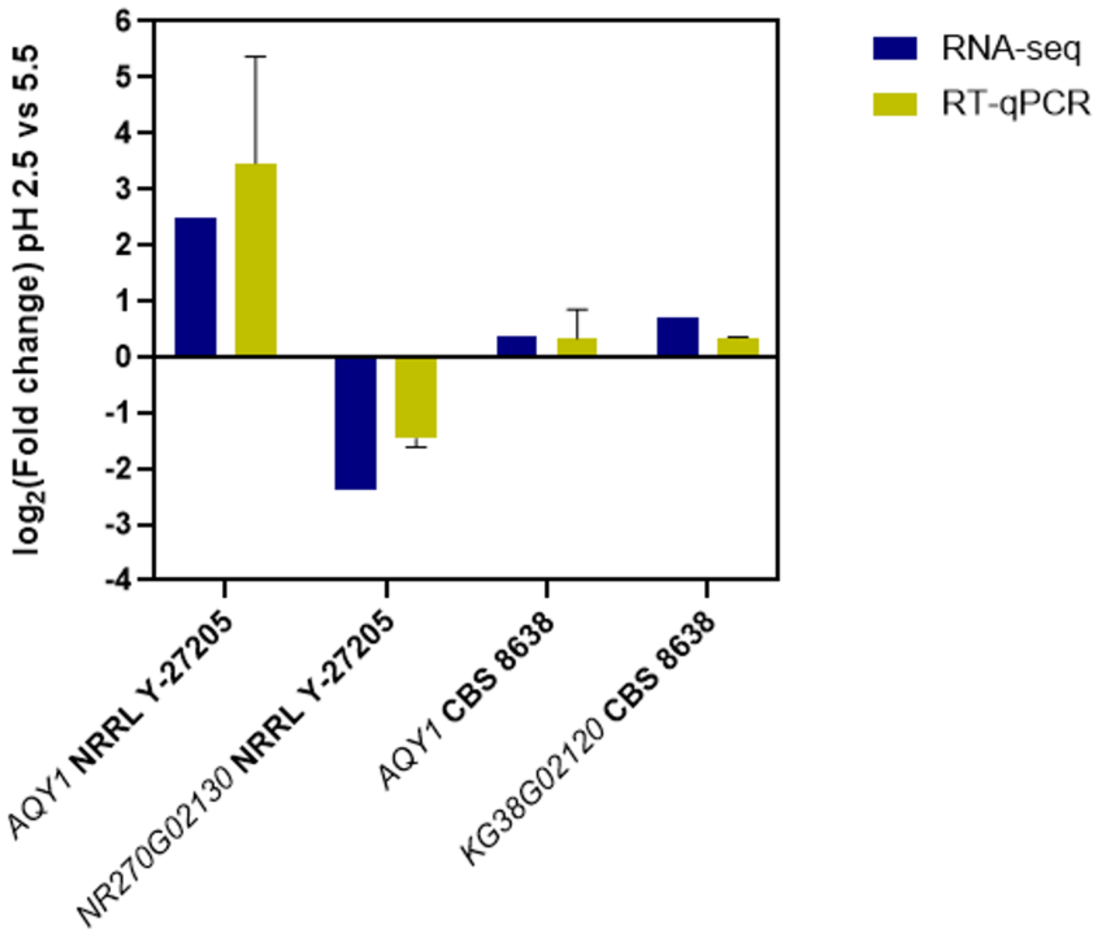

**Supplementary Figure 2. Enriched GO terms associated with the combined *M. bulderi* species response at pH 2.5 vs pH 5.5**

The plots show the following: Cellular component (CC) terms in upregulated genes (Panel A) and downregulated genes (Panel B); and molecular function (MF) terms in upregulated genes (Panel C) and downregulated genes (Panel D). The gene ratio value is the ratio between the DE genes over the total of DE genes in the same category and the dot size corresponds to the number of DE genes linked to that function as a fraction of total DE. The GO terms reported are those significantly enriched in DE genes (p-adj<0.01).

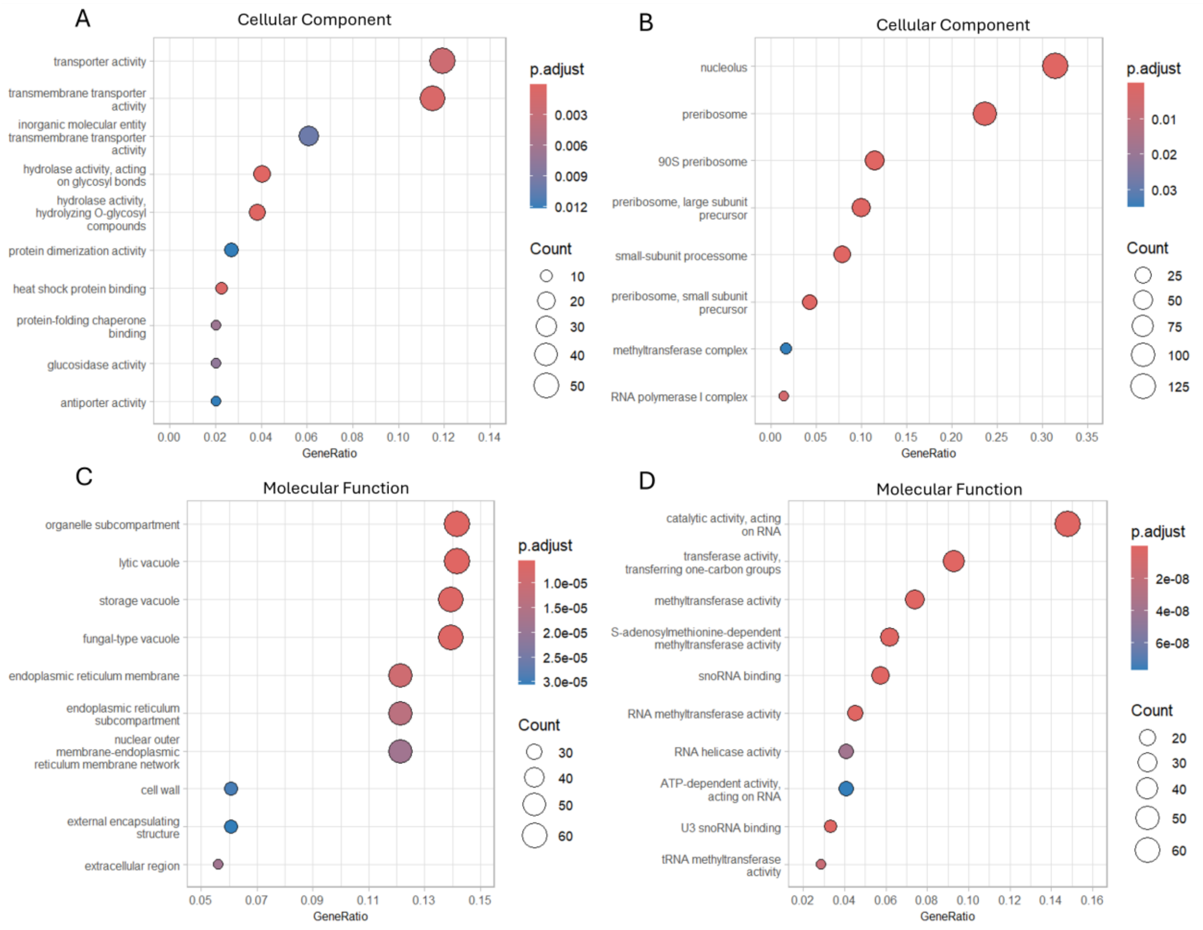

**Supplementary Figure 3. Enriched GO terms associated with the *M. bulderi* strain CBS8638 response at pH 2.5 vs pH 5.5**

The plots show the following: Biological process (BP) terms , in upregulated genes (Panel A) and downregulated genes (Panel B); CC terms in upregulated genes (Panel C) and downregulated genes (Panel D); and MF terms in upregulated genes (Panel E) and downregulated genes (Panel F). The gene ratio value is the ratio between the DE genes over the total of DE genes in the same category and the dot size corresponds to the number of DE genes linked to that function as a fraction of total DE. The GO terms reported are those significantly enriched in DE genes (p-adj<0.01).

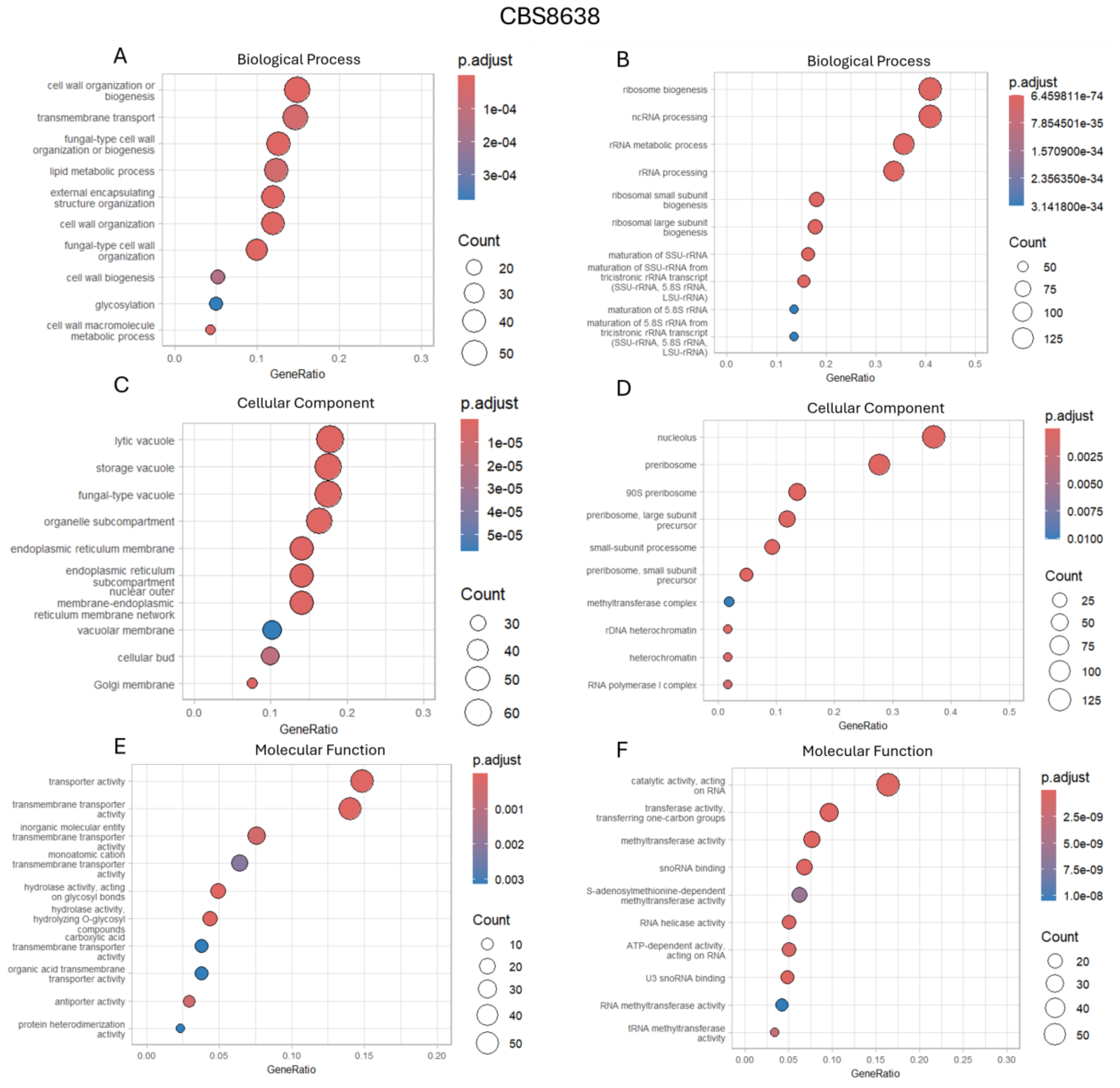

**Supplementary Figure 4. Enriched GO terms associated with the *M. bulderi* strain CBS86389 response at pH 2.5 vs pH 5.5**

The plots show the following: Biological process (BP) terms , in upregulated genes (Panel A) and downregulated genes (Panel B) and MF terms in upregulated genes (Panel C) and downregulated genes (Panel D). The gene ratio value is the ratio between the DE genes over the total of DE genes in the same category and the dot size corresponds to the number of DE genes linked to that function as a fraction of total DE. The GO terms reported are those significantly enriched in DE genes (p-adj<0.01). No GO terms associated with CC were significantly enriched in CBS863839.

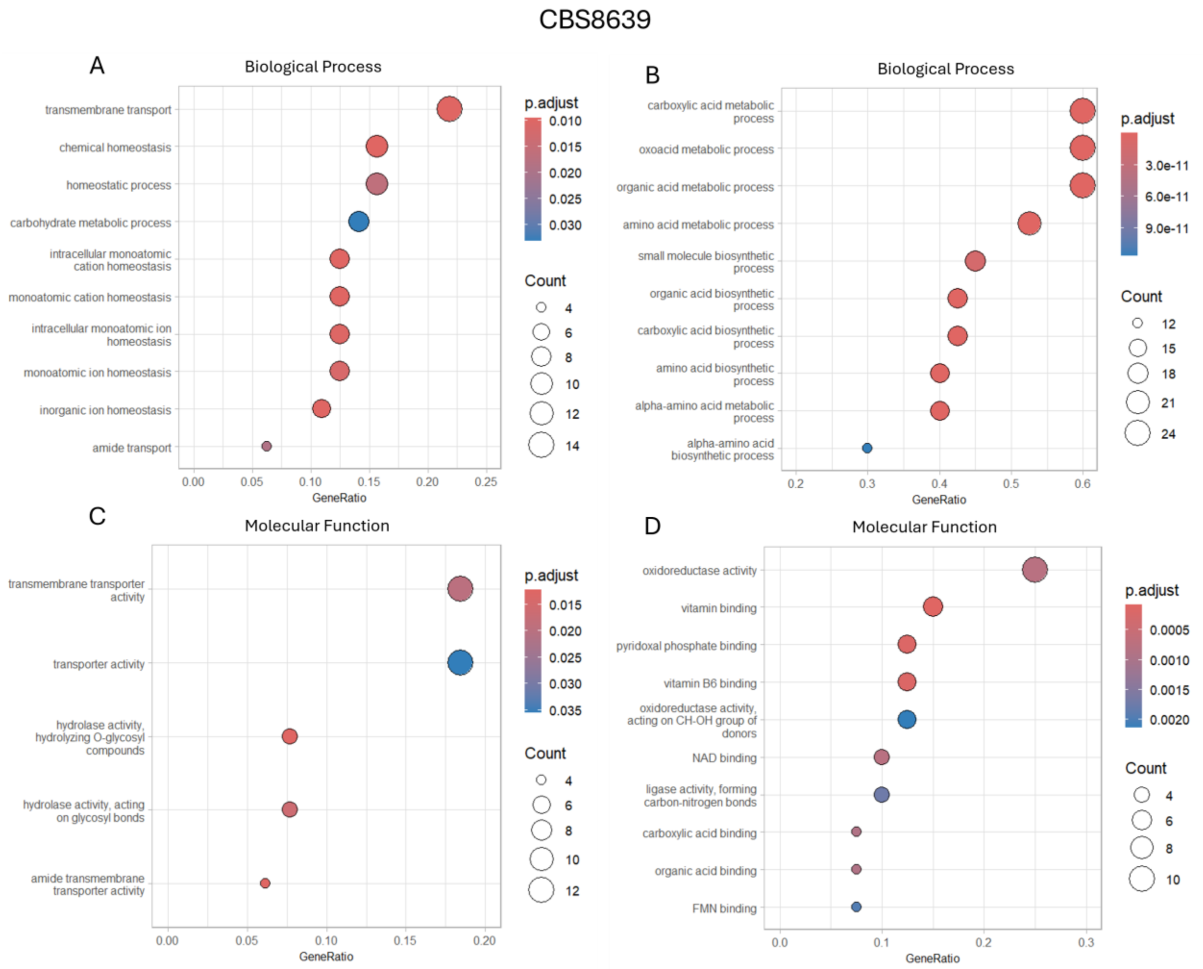

**Supplementary Figure 5. Enriched GO terms associated with the *M. bulderi* strain NRRL-Y27205 response at pH 2.5 vs pH 5.5**

The plots show the following: Biological process (BP) terms, in upregulated genes (Panel A) and downregulated genes (Panel B); CC terms in upregulated genes (Panel C) and downregulated genes (Panel D); and MF terms in upregulated genes (Panel E) and downregulated genes (Panel F). The gene ratio value is the ratio between the DE genes over the total of DE genes in the same category and the dot size corresponds to the number of DE genes linked to that function as a fraction of total DE. The GO terms reported are those significantly enriched in DE genes (p-adj<0.01).

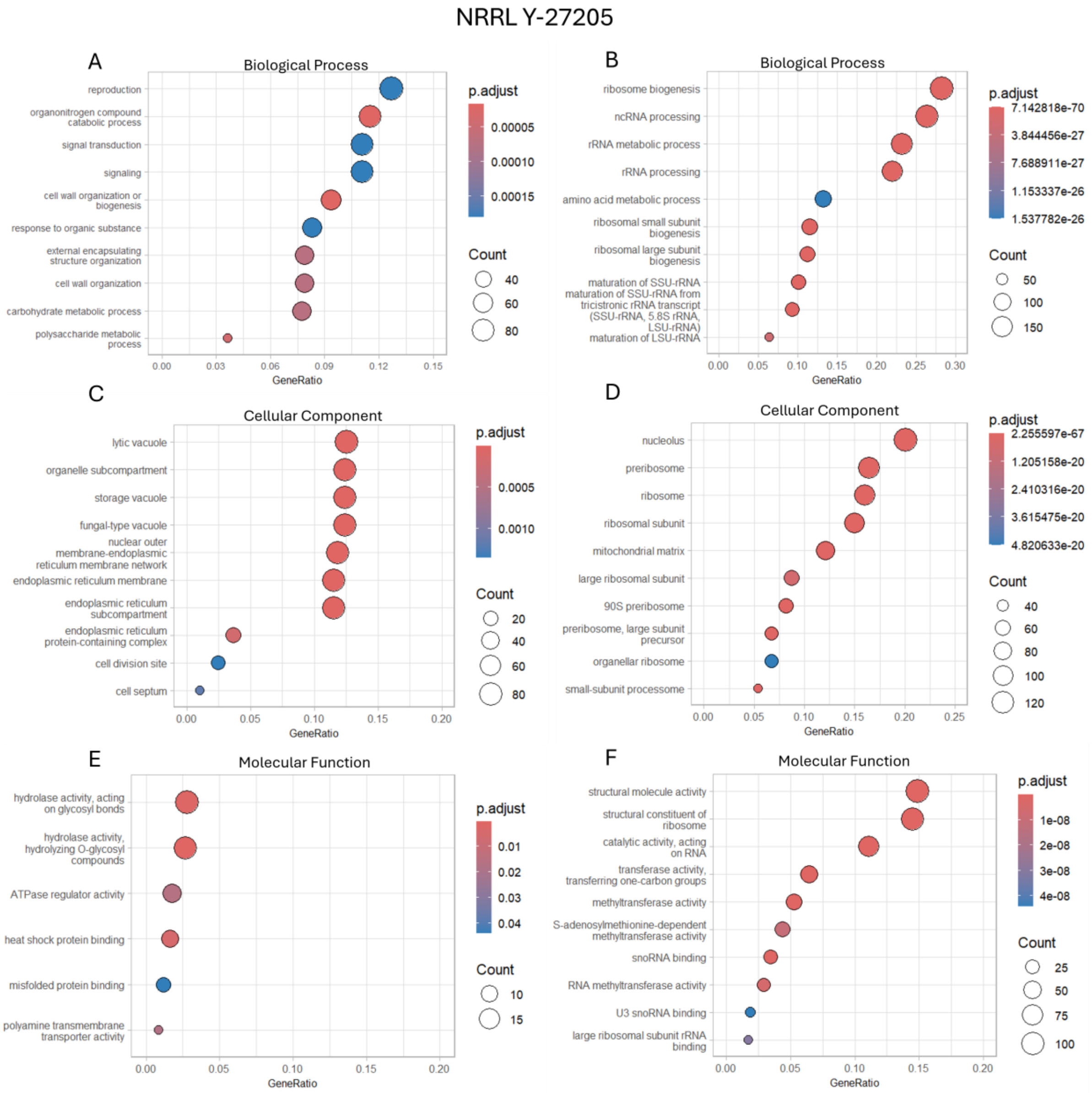

**Supplementary Figure 6. Cell wall morphology of *M. bulderi* and *S. cerevisiae* at different pHs**

Scanning electron micrographs of *M. bulderi* and *S. cerevisiae* grown in minimal media under the following conditions: Panel A: NCYC505 pH 5.5, Panel B: NCYC505 pH 2.5, Panel C: CBS8638 pH5.5, Panel D: CBS8638 pH 2.5, Panel E CBS8639 pH 5.5, Panel F: CBS8639 pH 2.5, Panel G: NRRL Y-27205 pH 5.5, Panel H: NRRL Y-27205 pH 2.5

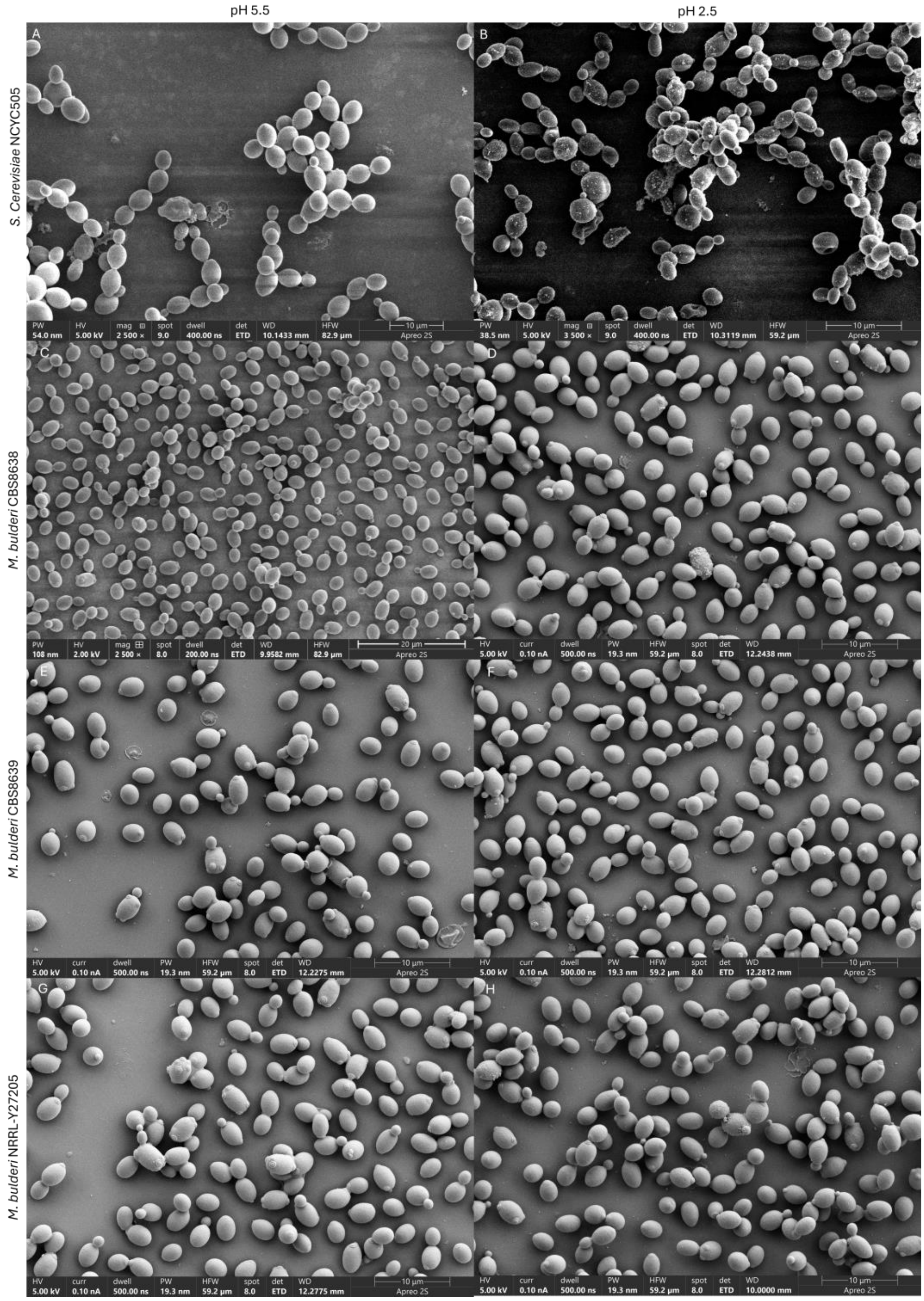

**Supplementary Figure 7. Analysis of lipid mass of *M. bulderi* and *S. cerevisiae***

Chart showing total lipid mass per gram of dried cell found in *S. cerevisiae* and *M. bulderi* grown at pH 2.5 or pH 5.5.

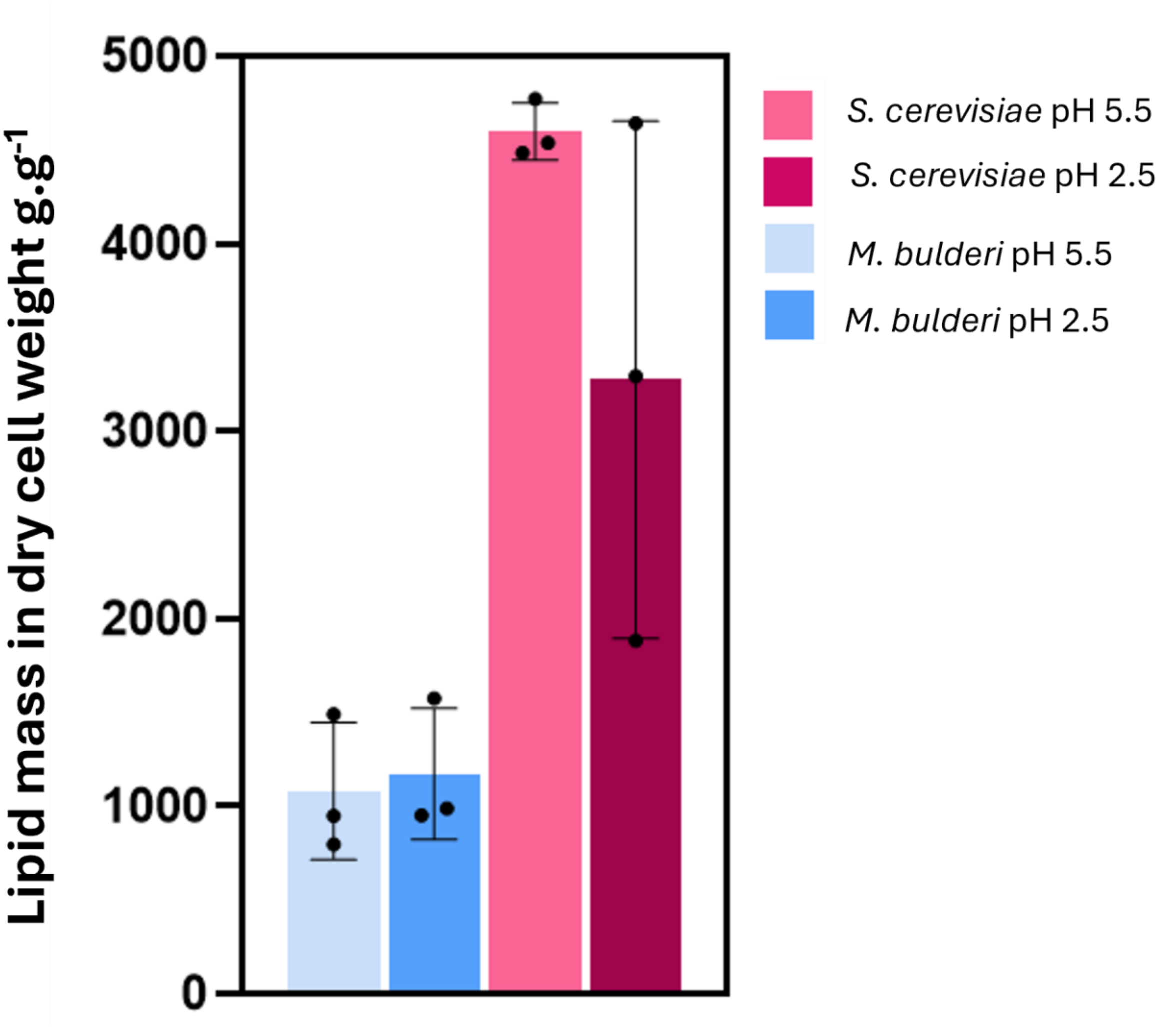
